# Gossypol biosynthesis in cotton revealed through organ culture, plant grafting and gene expression profiling

**DOI:** 10.1101/173138

**Authors:** Tianlun Zhao, Jiahui Hu, Cheng Li, Cong Li, Lei Mei, Jinhong Chen, Shuijin Zhu

## Abstract

Gossypol plays an important role in defense mechanism of *Gossypium* species and the presence of gossypol also limits the utilization of cottonseeds. However, little is known about the metabolism of gossypol in cotton plant. Here, Detection on the dynamic tendency of gossypol content illustrated that at the germination stage, the main source of gossypol was cotyledon, and at the later stages, gossypol mainly came from root system. Plant grafting between cottons and sunflower proved that gossypol was mainly synthesized in the root systems of cotton plants and both of the glanded and glandless cottons had the ability of gossypol biosynthesis. Besides, the pigment glands expression was uncoupled with gossypol biosynthesis. Root tip and rootless seedling organ culture in vitro further revealed other parts of the seedlings also got the ability to synthesize gossypol except root system. Moreover, root system produced the racemic gossypol and plant synthesized the optically active gossypol. The expression profiling of key genes in the gossypol biosynthetic pathway suggested that downstream key genes had relatively high expression levels in root systems which confirmed that gossypol was mainly synthesized in the root systems. Taken together, our results helped to clarify the complex mechanism of gossypol metabolism.

## Introduction

Cotton (*Gossypium spp.*) is one of the most important economic crops in the world. Cotton not only produces natural fiber for textile industry, but it also provides a large quantity of cottonseeds which contain high-quality protein and oil (Kong *et al.*, 2010). It is estimated that every kilogram fiber yield is coupled with 1.65 kg cottonseeds, which contain approximately 21% oil and 23% protein (Sunikumar *et al.*, 2006). However, cottonseeds cannot be used directly due to the presence of toxic gossypol and other related terpenoids, which are toxic to human beings and monogastric animals (Stipanovic *et al.*, 1975). On the other hand, the gossypol plays an important role in self-protection of cotton plants (Cai *et al.*, 2010; William *et al.*, 2011; Mellon *et al.*, 2012). Therefore, researchs on the gossypol metabolism are urgently needed in cotton production.

Gossypol was first characterized by Adams et al. in 1938 through a classic series of studies (Heinstein *et al.*, 1962). It is a polyphenolic aldehyde which constitutes 20-40% of the pigment glands weight and accounts for 0.4-1.7% of the whole cottonseed kernel. As a phytoalexin, gossypol provides constitutive and inducible resistance against the pest and pathogens (Carrière *et al.*, 2004; Wang *et al.*, 2004; Mao *et al.*, 2007; Stipanovic *et al.*, 2008). It was previously reported that gossypol-rich diet clearly reduced weight of bollworm larvae and affected their growth stages (Carrière *et al.*, 2004; Stipanovic *et al.*, 2008). Moreover, gossypol significantly inhibited the growth of filamentous fungi *Aspergillus flavus* (Mellon *et al.*, 2012). Besides pest control, gossypol can be used as anti-cancer (Cotyle *et al.*, 1994; Liu *et al.*, 2002; Ye *et al.*, 2007), anti-bacterial (Radloff *et al.*, 1986; Tegos *et al.*, 2002) and male contraceptive (Wang *et al.*, 1987; Coutinho, 2002; Lopez *et al.*, 2005). There are two different enantiomers of gossypol, (-)-gossypol and (+)-gossypol. Based on previous studies, the biological activity of (-)-gossypol is stronger than (+)-gossypol (Puckhaber *et al.*, 2002; Wolter *et al.*, 2006; Kline *et al.*, 2008; Mellon *et al.*, 2011). Futher, the differences in ratio of enantiomers in cottonseeds might affect the poultry production when they were used as poultry feed (Bailey *et al.*, 2000).

Recently, several key genes involved in the pathway of the gossypol biosynthesis have been identified and characterized. Two terpene synthase genes, *GhTPS1* and *GhTPS2* (Huang *et al.*, 2013), and two genes which were the limiting enzyme genes in the pathway of isoprenoid biosynthesis, *hmg1* and *hmg2* (Leandro *et al.*, 1999) have been elucidated. Other key enzymes in the pathway of gossypol biosynthesis whose encoding genes exist as gene families, such as *CAD1*-*A*, *CAD1*-*C2* (Meng *et al.*, 1999) and *cdn1*-*C4* (Townsend *et al.*, 2005) have been identified and researched. Similarly, *GaCYP706B1* was isolated as the encoding gene of cadinene-8-hydroxylase which is the key enzyme in the hemigossypol biosynthesis (Luo *et al.*, 2001). Besides, a transcription factor, *GaWRKY1*, might affect the expression of *CAD*-*1* to regulate the biosynthesis of sesquiterpene in cotton (*Xu et al.*, 2004). Nowadays, as tremendous progress in molecular biology has been made, gene transformation, virus-induced gene silencing (VIGS) and CRISPR/Cas9 system have been applied to study cotton traits (Sunikumar *et al.*, 2006; Ma *et al.*, 2016; Li *et al.*, 2017), which provides a good opportunity to figure out the complex mechanism of gossypol metabolism by combining genetic and physiological approaches.

Several studies have attempted to explain the relationship between pigment gland and gossypol content in cotton plants. Punit et al. found out that different genotypes had an impact on the distribution of pigment glands (Punit *et al.*, 1991). The gossypol content in cotton was closely related to the genetic types of pigment glands in cotton cultivars. Singh and Weaver proposed that the gossypol content was highly correlated with the number of pigment glands (Punit and Singh, 1972), except *G. somalense* Huntch, which had almost no gossypol in seed although normal pigment glands are existed (Xiang and Yang, 1993). Silencing the *CYP706B1*, a key gene in gossypol biosynthesis pathway, Ma et al. found that the gossypol content was significantly reduced but the pigment glands, as in a normal plant, were still formed, while eliminating the pigment glands through VIGS led to decline in gossypol content greatly (Ma *et al.*, 2016). Therefore, the relationship between gossypol and pigment glands was complicated, which needs more evidence to clarify.

Understanding the metabolism of gossypol could be conductive to develop lower gossypol seed cotton cultivars that could be well utilized and widely adapted.

However, only few studies have been carried out to elucidate the biosynthesis and transportation of gossypol physiologically. Smith proposed that the gossypol was synthesized in cotton root via root culture in vitro (Smith, 1961). However, no report had proved whether other tissues had the ability to synthesize gossypol or not. To clearly define the gossypol biosynthesis and transportation in cotton plants is not only useful for understanding the mechanism of gossypol biosynthesis theoretically, but ultimate goal is helping the scientists to develop the new cotton cultivars with lower gossypol content in seeds by genetic engineering.

In present study, plants grafting, organ culture and gene expression profiling were used to figure out gossypol biosynthesis and its transportation, and to clarify the relationship between gossypol and pigment glands in cotton plant, which may be helpful to elucidate the complicated mechanism of gossypol metabolism and the key roles of gossypol in cotton ontogeny.

## Materials and methods

### Plant material and sampling

Two pairs of glanded and glandless upland cotton isogenic lines, CRI17 and CRI17W, Coker 312 and Coker 312W (developed in Zhejiang University), were used in the experiments. Other upland cotton materials, CRI49, Z5629 and TM-1, used in the experiment were as follows: CRI49, an extending traditional cotton cultivar of China, was provided by Cotton Research Institute; Z5629, a glandless germplasm derived from Zhejiang University; TM-1, the standard upland cotton line widely used in cotton research, was provided by USDA-ARS, College Station, Texas, USA. All the materials were kept by selfing in Zhejiang University.

Sunflower (*Helianthus annuus* L.) cultivar Sandaomei (SDM) was used as the rootstock material in this study. This was obtained from Jilin Province, China.

### Plant grafting

CRI49, CRI17W, Z5629 and SDM were grown in the greenhouse of Zhejiang University, China. The seedlings with two true leaves were used as the scions and rootstocks. The glanded cotton scions, CRI49, were grafted on glandless cotton rootstocks (CRI17W and Z5629) and sunflower rootstocks (SDM) by the method of improved bark graftage. Likewise, the glandless cotton scions, CRI17W and Z5629 were also grafted on glanded cotton rootstocks (CRI49).

After the operation of grafting, the soil was watered, and the grafted cotton seedlings were covered with the plastic bags and grown under 28 ± 2 . About four days later, the seedlings were covered by the plastic bags poked with 4 small holes for gas exchange. Two weeks later, the plastic bags were removed from the seedlings.

Leaves from the grafted cotton were sampled during the growth season and cottonseeds were sampled at the harvest stage for determination of gossypol content in three biological replications. All collected samples were immediately frozen in liquid nitrogen and stored at −80 °C.

### Tissue and organ culture in vitro

Cottonseed kernels were first washed by sterile water for 3 min twice, then disinfected by 70% alcohol for 3 min twice, followed by sterilization in mercuric chloride for 10 min, and finally washed for 2 min in sterile water five times. The kernels were germinated on the MS medium. When seed root attained to the length of 2 cm, root tip (about 1cm) was cut from the seedling for root tip culture in vitro, and the seedlings without root were used for culture in vitro as well.

The medium for root tip culture in vitro was that of macro nutrients and 1/2 micro nutrients of MS basic medium, vitamins and organic materials of B6 medium containing 100 mg/L of inositol, 1 mg/L of nicotinic acid, 10 mg/L of vitamin B1, 1 mg/L of vitamin B6, 20 mg/L of sucrose and 0.125 mg/L of IBA. Using 0.8% of agar as the solid material, and the medium was adjusted to pH to 6.4. Growth conditions for root tip culture were 14/10 h day night intervals with light intensity of 2000 lux and average temperature of 28±2 °C.

Rootless seedling culture in vitro was similar to root tip culture, except slight change in the medium, in which the MS basic medium, vitamins and organic materials of B6 basic medium were used with 0.1mg/L kinetin, 20 mg/L of sucrose. Also 0.8% of agar was used as the solid material and pH was adjusted to 6.4.

### Gossypol determination

Gossypol enantiomers were determined by HPLC. Standard gossypol solutions were prepared by dissolving 0.01 mg of HPLC-grade gossypol in 10 mL of acetonitrile and then adding 0.001, 0.002, 0.005, 0.050, 0.100, 0.200, 0.500, 0.800, 1.000, 2.000 and 3.000 mL into 2 mL derivative reagents which contained 2% D-alaninol, 10% acetic acid and 88% acetonitrile.

All samples were dried at 30 °C to constant weight for 3-4 days, and then ground with grinder to powder, and stored at −80 °C. 0.10 g of each sample was suspended into 2 mL derivative reagent to form sample solution.

Standard gossypol and sample solutions were water bathed at 75  for 45 min. Sample suspensions were filtered through quantitative filter paper followed by a filtration with a 0.45 μm syringe filter (Agela, Newark, USA). The sediment was washed three times by acetonitrile. After this procedure, the extract was adjusted to 10 mL using acetonitrile.

HPLC analysis were performed on Agilent 1100 (Agilent, Santa Clara, USA), equipped with an auto-sampler and an UV detection. A C18 column (250 mm × 4.6 mm, 5 μm, Dikma, Richmond Hill, USA) was employed as stationary phase. The mobile phase consisted of acetonitrile/0.2% H_3_PO_4_ (75/25, v/v). Injection volume was 10 μL and the flow rate was 1.0 mL/min. The UV detector was set at 238 nm and the temperature was 25 °C. Samples were measured in triplicates.

### Image analysis

The pigment glands of the leaves were observed and taken as images through an Olympus dissecting microscope (LEICA MZ95, Germany) with a digital camera. Density and size of the pigment glands in the images were measured by the Image Pro Plus (V6.0) software.

### Quantitative RT-PCR

True leaves, cotyledons, seed roots, secondary roots and stems were sampled from plant materials used in the experiment with three biological replicates. RNA of each sample was extracted using a Total RNA Extraction kit (Aidlab, Beijing, China). First strand cDNA was synthesized using TransScript One-Step gDNA Removal and cDNA Synthesis SuperMix (TransGen Biotec Co.,Ltd.) following manufactures protocol. Primers for quantitative RT-PCR (qRT-PCT) were designed with the Primer 5.0 software. All primers used in the experiment are listed in Supplementary Table S1. The amplification reactions of qRT-PCR were performed with Lightcycler 96 system (Roche) using SYBR the premix Ex taq (TakaRa) with following parameters: 30 s initializing denaturation at 95 °C; following 45 cycles of 10 s denaturation at 95 °C, 10 s annealing at 54 °C, and 20 s extension at 72 °C. In addition, the default setting for the melting curve stage was chosen. The relative expression levels were calculated by the method of 2^−ΔΔCt^. The heatmap for expression profiles was generated with the Mev 4.0 software.

### Statistical analysis

Statistical analysis for (±)-gossypol content and pigment glands was carried out by SPSS20.0. Data were represented as mean ± SD, *P* < 0.05, and values of *P* < 0.01 were considered as statistically significant and extremely significant.

## Results

### Changes in gossypol content of glanded and glandless cotton seedlings

Two pairs of NILS were grown in the greenhouse, and different parts of plants were sampled at different seedlings stages, and the contents of gossypol were detected (Fig. 1).

**Fig. 1.**
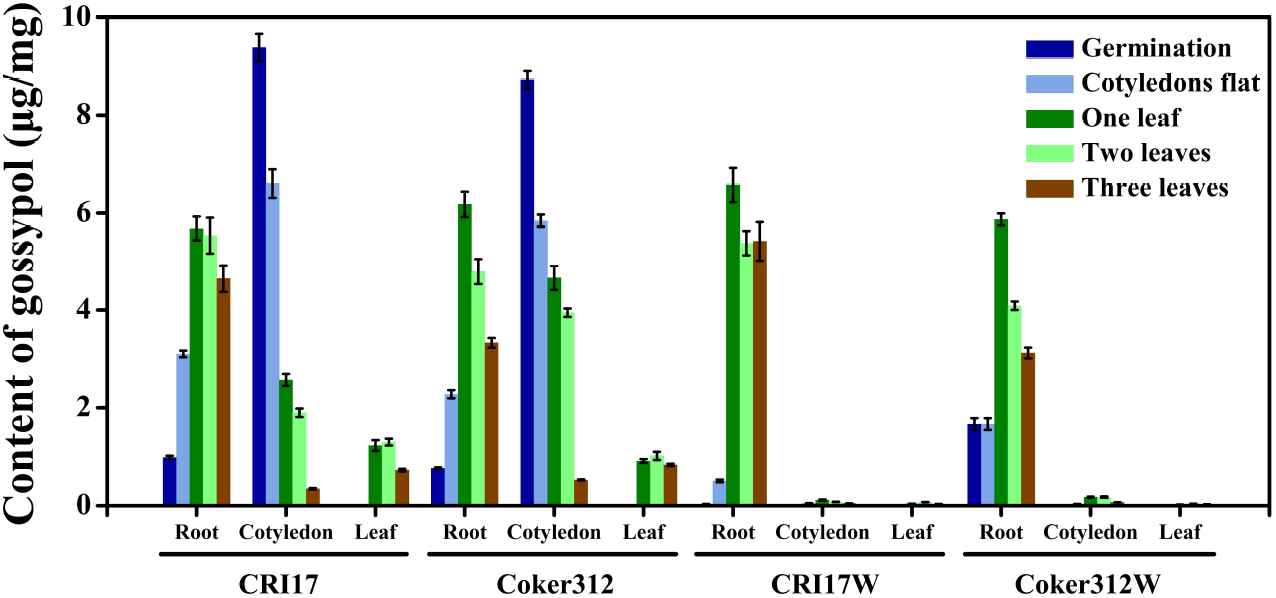
The dynamic changes of gossypol content in tissues of the cotton plants (glanded, dominant glandless and recessive glandless) at the germination and seedling stages.

In cotyledon, at the germination stage, the gossypol contents in glanded cotton, CRI17 and Coker312 (9.384 and 8.722 μg/mg), were much higher than that of their corresponding glandless isogenic lines, CRI17W and Coker312W (0.011 and 0.008 μg/mg). At the subsequent stages, gossypol content in CRI17 and Coker312 was decreased all the time. At three true leaves stage, their gossypol contents were reduced to 0.345 and 0.528 μg/mg. However, the gossypol contents increased and reached a peak at the one leaf stage in CRI17W (0.116 μg/mg) and two leaves stage in Coker312W (0.177 μg/mg). It was indicated that gossypol was either decomposed or transported from the cotyledons to other tissues in glanded cotton, while for their isogenic glandless plants, it was accumulated in cotyledon.

In seed roots, at the germination stage, the gossypol contents of CRI17 and Coker312 were 0.986 and 0.768 μg/mg, which were much higher than that in their glandless isogenic lines, CRI17W and Coker312W (0.033 and 0.023 μg/mg). As seedlings grew, the gossypol contents were increased and reached the peak levels at the one true leaf stage in all four cultivars. Moreover, the content in glandless plants was greater than that of their glanded isogenic lines. Thus, it was illustrated that the plant started to produce gossypol after seed germination,.

Similarly, in leaves, the gossypol contents of glanded cotton were much higher than their glandless isogenic lines. Although gossypol contents were increasing with the growth of gossypol and reached the peak at the two leaves stage. Exactly, the contents of gossypol at that stage were 0.130 and 0.102 μg/mg in CRI17 and Coker312, 0.007 and 0.004 μg/mg in CRI17W and Coker312W.

The dynamic tendency of gossypol suggested that the gossypol was synthesized in the root of glanded and glandless plants, and both of them possessed a strong ability to synthesize the gossypol. At germination stage, the main source of gossypol was cotyledon and from where it transported to the other parts. The roots system started to synthesize the gossypol at the cotyledons flat stage. At later stages, the gossypol was mainly synthesized in roots and transferred to other tissues.

### Pigment glands and gossypol changes after grafting

To investigate the effect of root system on the gossypol biosynthesis, different combinations of plant grafting were performed (Supplementary Fig. S2).

All combinations of grafted plants were developed as normal plants from which cottonseed were harvested. However, among the combinations of cotton scions and sunflower rootstocks grew very week and died after two weeks, except plants with regenerated roots which were derived from the scion stems grew very well for one month.

### Grafting between glanded and glandless cottons

The size and density of pigment glands in leaves and seeds were measured and compared between grafted plant and normal plants (Supplementary Table S2). Results indicated that there was no significant difference in the size and density of pigment glands between the normal glanded plant and the grafted glanded plant scions with glandless rootstock. Similarly, no pigment gland on the glandless cotton; either on normal plant or grafted plant with glanded rootstocks were observed.

Gossypol contents in cottonseeds of plants and grafted plants were highly varied. It was shown that when dominant glandless cotton, CRI17W was used as the rootstock, the gossypol contents in the cottonseeds of scion were significantly lower than that of the normal CRI49 plants (Fig. 2). In exact terms, the (+), (-) and (±)-gossypol were decreased by 23.24%, 28.14% and 25.14% respectively. In case of recessive glandless cotton, Z5629, used as the rootstock, the (+), (-) and (±)-gossypol decreased by 26.33%, 15.29% and 22.25% respectively. On the contrary, when the CRI49 was used as the rootstock, the gossypol contents in the seeds of glandless cotton scions were extremely significantly higher than their respective normal plants (Fig. 2). Accurately, for CRI17W, the (+), (-) and (±)-gossypol contents got 4.10, 3.76 and 4.06 times increase, and for Z5629, 2.45, 2.95 and 2.67 times respectively, as compared with their normal plants.

**Fig. 2.**
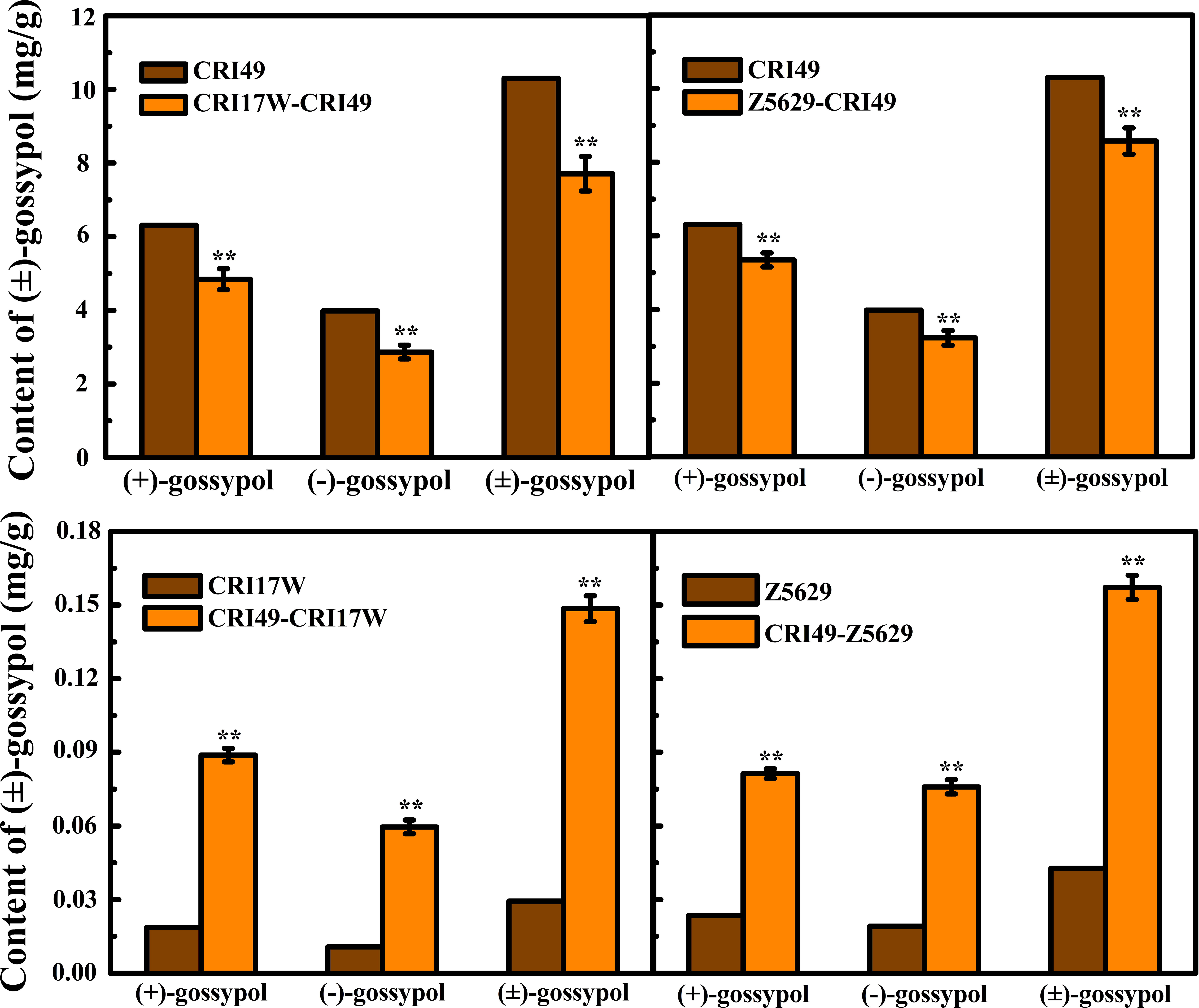
Changes of gossypol content in scions before and after grafting, ^∗∗^Significant at *P* = 0.01.

This suggested that root system impacted the content of gossypol in the aboveground part, but there was no effect on the expression of pigment glands. Furthermore, root systems of both glanded and glandless plants had the ability to synthesize (±)-gossypol, but their ability was differed.

### Grafting between sunflower and glanded cotton

To further confirm gossypol biosynthesis of cotton root system, the glanded cotton plants, CRI49, were grafted on the sunflower rootstocks which are unable to synthesize gossypol. Due to species isolation, the scions only survived for around half a month; however, scions with the regenerated roots derived from the bottom of the scions survived for a month (Fig. 3).

**Fig. 3.**
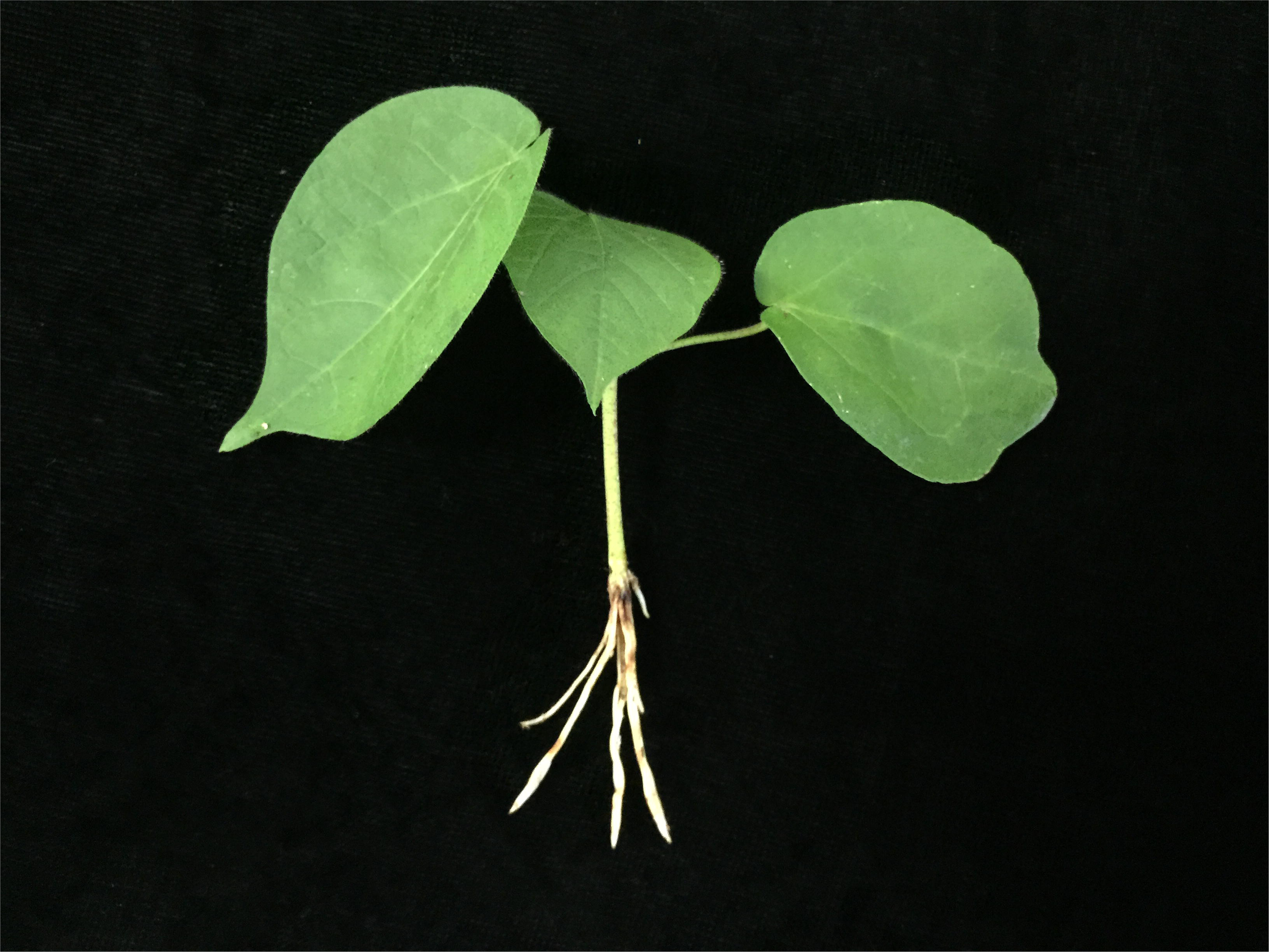
Cotton scions with the generated roots around eight days after grafting on the rootstocks of sunflower.

The dynamic content of (±)-gossypol in scion leaves was detected by HPLC method. As the results shown (Fig. 4), at 0 day of grafting, the contents of (+), (-) and (±)-gossypol in scion leaves were 0.364, 0.320 and 0.684 μg/mg, respectively. Four days after grafting, the contents of (-)-gossypol and (±)-gossypol in the scion leaves declined by 19.53% and 14.58% as compared with 0 days, and (+)-gossypol was significantly decreased by 10.23%. Twelve days after grafting, compared with eight days, the contents of (+), (-) and (±)-gossypol in the scions without regenerated root still got extreme significant decrease by 16.64%, 33.28% and 23.11% respectively. Meanwhile, the contents of (+) and (±)-gossypol in the scions with regenerated roots were still increased by 5.37% and 8.48%, but they were not significant statistically, while the content of (-)-gossypol showed a significant increase by 12.21%. On the other hand, compared with the scions without regenerated roots, the content of (+), (-) and (±)-gossypol in the scions with regenerated roots were extremely and significantly higher at the same time after grafting. Thus, it was indicated that regenerated root had the ability to synthesize gossypol.

**Fig. 4.**
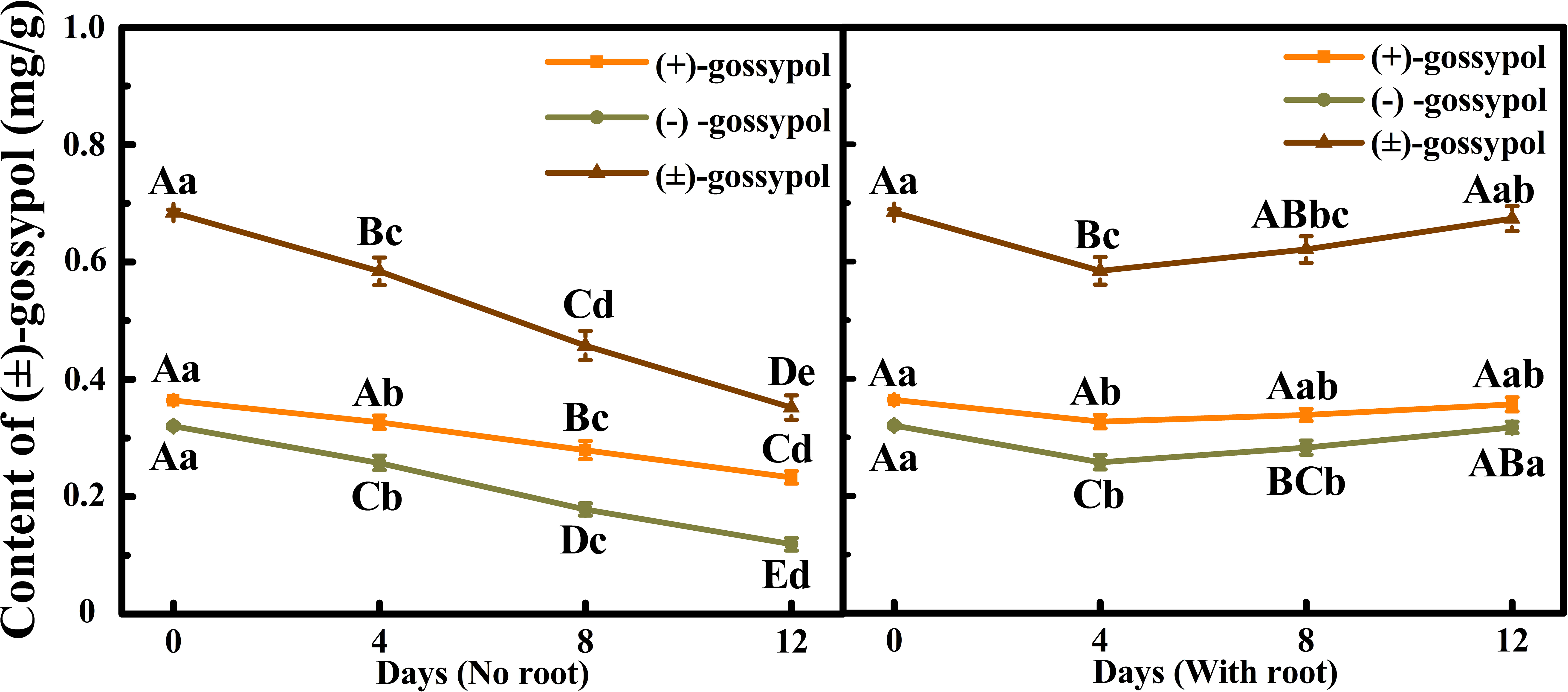
Dynamic changes of (+), (-) and (±)-gossypol content in CRI49 leaves before and after grafting on the rootstock of sunflower. Different letters indicate significant difference at the different times. Uppercase and lowercase letters indicate significant at *P*=0.01 and 0.05 respectively.

### Gossypol change during the tissues culture in vitro

In order to investigate the metabolism of gossypol, the root tip culture and rootless seedlings (plant without root tip) culture in vitro were performed (Supplementary Fig. S2). Both root system and the rootless seedling grew well in the media. The gossypol contents in the root system and rootless seedlings were measured during the growth period.

Results showed that the gossypol contents in the root systems of the glanded cultivars, CRI17 and Coker312, were 0.986 and 0.768 μg/mg at 0 day, respectively (Fig. 5). Two days after incubation, these plants showed extremely significant decrease by 46.06% and 32.22%, and then a continuous extremely significant increase occurred afterwards. At the 16 days of culture, the gossypol contents were 3.72 and 4.22 times compared to those of 0 day. Similarly, for the root tip culture in the glandless cotton, CRI17W and Coker312W, the gossypol contents were 0.033 and 0.023 μg/mg only at 0 day of culture, but were increasing all the time during culture period. At the 16 days of culture, the gossypol contents reached to 134.47 and 101.42 times compared to those of 0 day. Comparing the gossypol contents between glanded and glandless cotton root systems during cultured in vitro, it was clearly indicated that both had almost same ability of gossypol biosynthesis, although there was a significant difference in gossypol content at two days which was due to the initial gossypol contents of the cottonseeds.

**Fig. 5.**
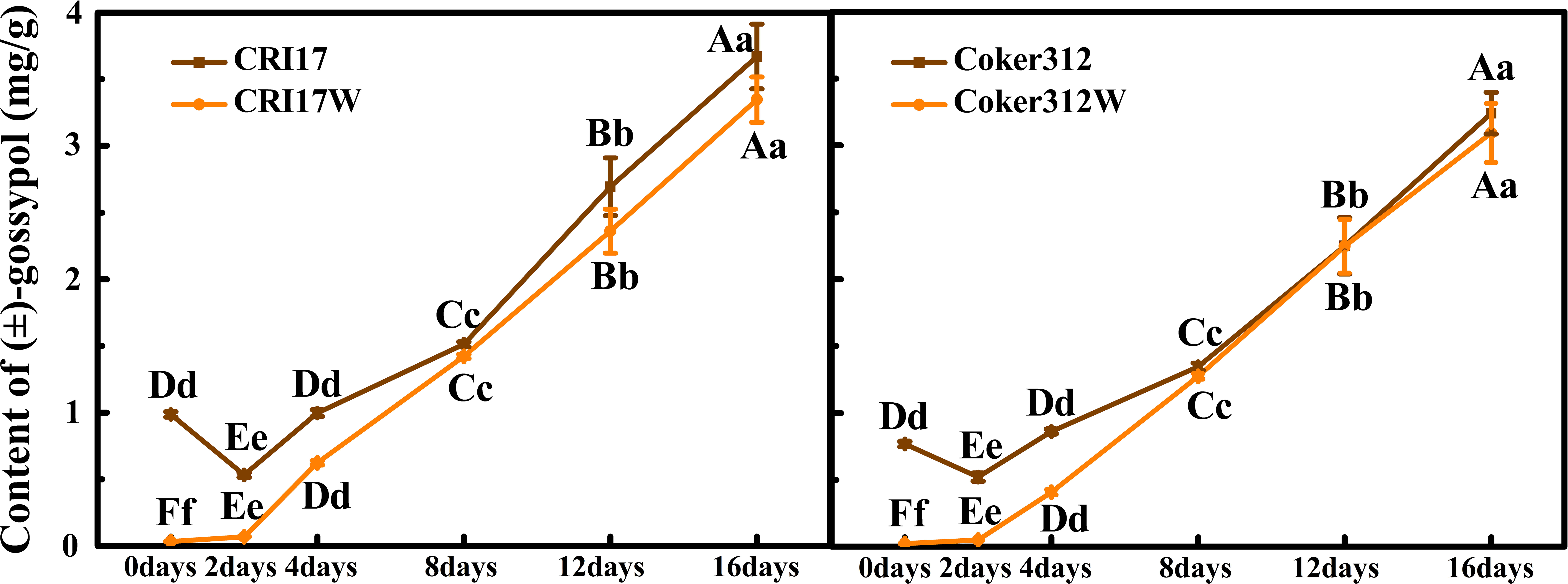
The dynamic changes of gossypol content in the root systems of four cultivars (glanded, dominant glandless and recessive glandless). Different letters indicate significant difference at the different times. Uppercase and lowercase letters indicate significant at *P*=0.01 and 0.05 respectively.

The rootless seedlings culture in vitro produced two types of products, rootless seedlings and regenerated root seedlings. The dynamic changes of gossypol contents are shown (Fig. 6). For the glanded seedlings, both with and without generated roots, the gossypol contents were decreased all the time due to presence of high gossypol content in seeds. However, the gossypol contents of the seedlings with the regenerated roots were extreme significantly higher than that of the seedlings without the generated root. Comparing to the four days after incubating, the gossypol contents in CRI17 and Coker312 with the regenerated roots decreased to 58.06% (3.924 μg/mg) and 49.39% (33.14 μg/mg) at 16 days after culture, while those without the regenerated roots decreased to 25.51% (1.724 μg/mg) and 23.67% (1.588 μg/mg) only, respectively. On the contrary, in the glandless plants with or without regenerated roots, the gossypol contents had been increasing, and contents in the seedlings with the regenerated root were extremely significant higher than the rootless seedlings. Futher it was observed that at eight days after culture, the gossypol contents in CRI17W and Coker312W with the regenerated root increased to 2.47 times (0.833 μg/mg) and 3.02 (0.858 μg/mg) times increase at the 16 days, while the seedlings without the root increased to 1.40 times (0.471 μg/mg) and 1.60 times (0.454 μg/mg), respectively. By comparing the gossypol contents between glanded and glandless rootless seedlings cultured in vitro, it was figured that the gossypol contents in glanded seedlings were extreme significantly higher than that of glandless ones at the initial stage. Then the gossypol contents decreased in glanded seedlings, while increased in glanded seedlings. It was implied that the growth and development of seedlings need a certain amount of gossypol, which is not enough in the glandless cotton, but too much for glanded cotton in vitro. So the rootless glandless cotton seedlings must produce gossypol for their growth and development, which also approved at the same time that the cotton plant can synthesize gossypol as well.

**Fig. 6.**
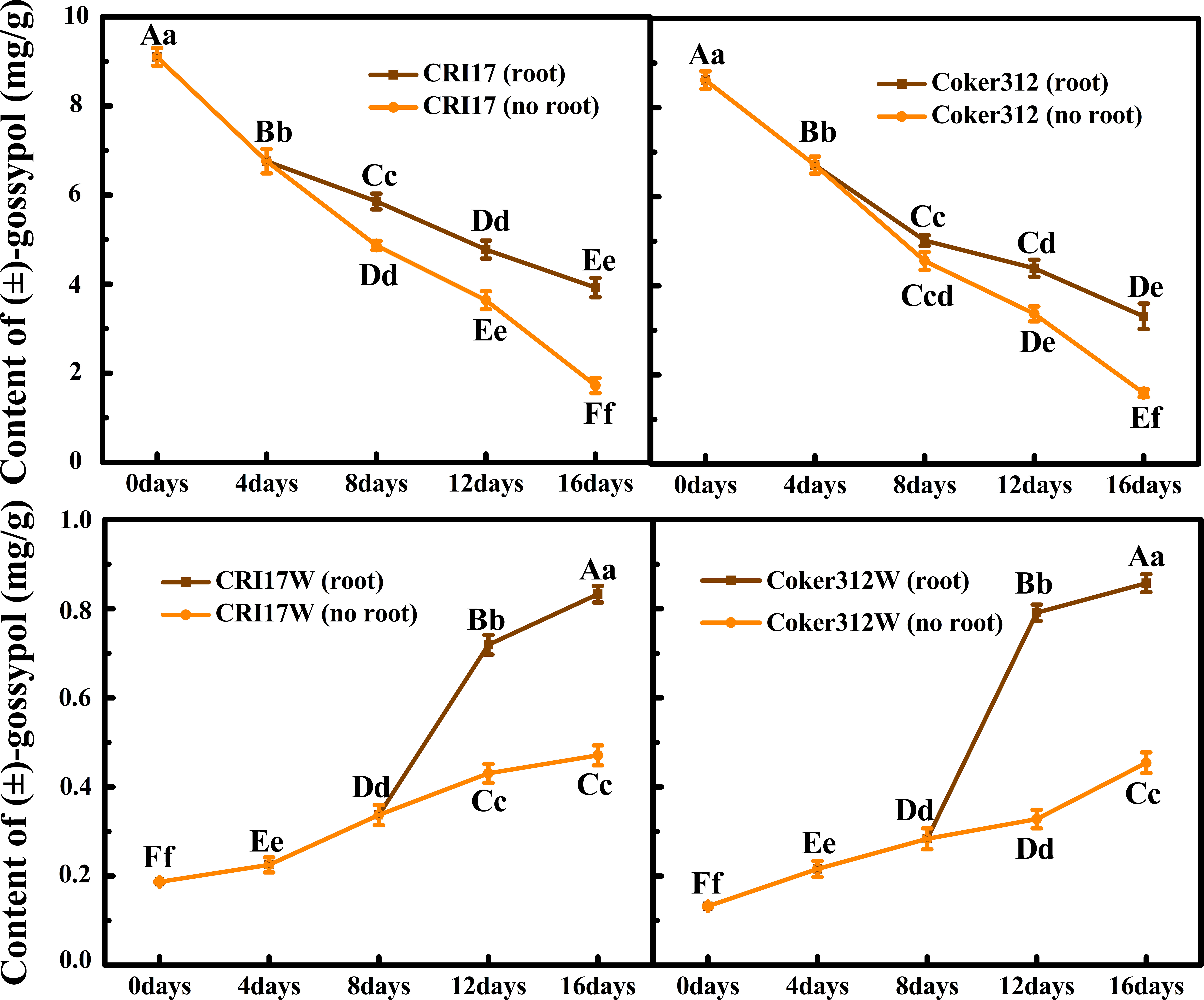
The dynamic changes of gossypol content in the rootless seedlings (with and without generated root) of four cultivars (glanded, dominant glandless and recessive glandless). Different letters indicate significant difference at the different times. Uppercase and lowercase letters indicate significant at *P*=0.05 and 0.01 respectively.

To investigate the gossypol isomers produced from different organs at different stages, the content of (+) and (-)-gossypol during the root tip culture and the rootless seedlings culture in vitro were measured (Supplementary Table S2 and Supplementary Table S3). The ratio of (+)/(-)-gossypol in the process of root tip culture maintained stable around one (Fig. 7), while in the rootless seedlings, the ratio was more than one in the seedlings with the regenerated root for all the times after the regenerated roots emerged. It implied that the gossypol produced by root system was racemic gossypol, which was different from that produced by plant in which the (+)-gossypol was more than (-)-gossypol. Besides, it was illustrated that the ability of root system to synthesize gossypol was much greater than other organs.

**Fig. 7.**
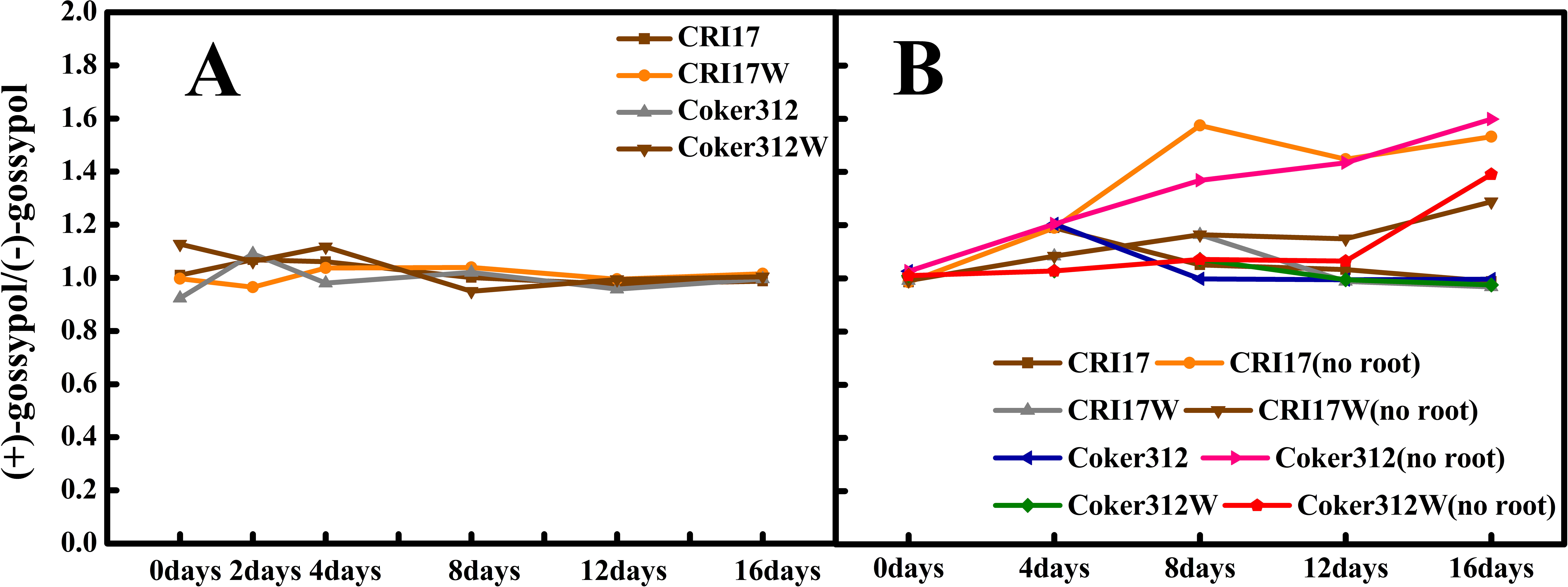
The ratio of (+)/(-)-gossypol in the process of the organs culture. (A) Root tip culture. (B) Rootless plant culture.

### Expression patterns of the key genes in the gossypol biosynthetic pathway

Several key genes in the biosynthesis pathway of gossypol were selected and quantitative RT-PCR analysis in true leaf, cotyledon, secondary root, seed root and stem of CRI17, CRI17W, Coker312, Coker312W and TM-1 was taken to reveal the expression patterns of these key genes.

As the results showed (Fig. 8), the key genes related to gossypol biosynthesis all had their specific expression patterns. In the upstream of the gossypol biosynthesis pathway, HMGR is considered one of the most important key enzymes. The corresponding encoding genes, *hmg1* and *hmg2*, both were highly expressed in root systems of five materials. Especially for *hmg1*, the expression level in seed root was highest. For the next key gene selected, *FPS*, the expression level in the root system was highest, followed by true leaves. *CAD1*-*A* is another key gene in gossypol biosynthesis pathway. It was more highly expressed in roots system of all cultivars than other tissues. Moreover, the expression level of *CAD1*-*A* in seed root was higher than that in secondary root. In the downstream of gossypol biosynthesis pathway, key gene *CYP706B1* also got the highest expression level in root system. In addition, *WRKY1*, a transcription factor in gossypol biosynthesis, demonstrated high expression levels in cotyledon and stem, and followed by root system. So, it is revealed that all the key genes in gossypol pathway biosynthesis had a relative high expression level in the root system except *WRKY1* and a relative low expression level in the leaves. The later genes participating in the gossypol biosynthesis pathway, the expression levels were higher in root systems. Therefore, it suggested that the root system was the main organ responsible for gossypol biosynthesis in cotton plant.

**Fig. 8.**
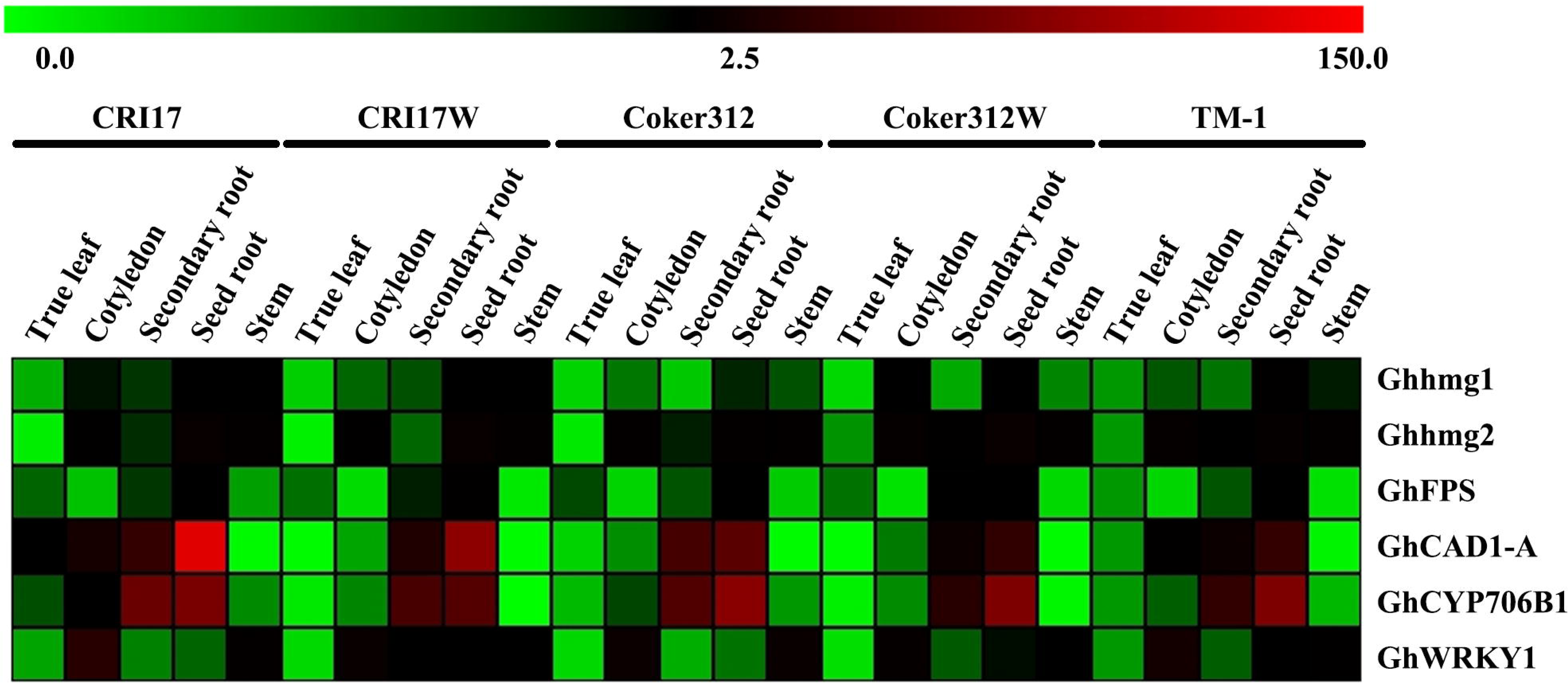
Expression patterns of the key genes participating in the pathway of gossypol biosynthesis. The color scales represents the relative signal intensity values.

## Discussion

Gossypol is one of the most important secondary metabolites for cotton plant growth and development (Adams *et al.*, 1960; Zhou *et al.*, 2013). In our result that the glanded rootless seedlings which had more gossypol generated the adventitious roots significantly earlier than that of glandless ones, which might imply that the gossypol have an effect on root regeneration and it also proved the importance of gossypol for plant development. However, the mechanism of gossypol biosynthesis and transportation are still poorly known. Previously, Smith proposed that gossypol was synthesized in the excised root by root culture in vitro (Smith, 1961), but he had not proved whether other plant organs had the ability to synthesize the gossypol or not. In this study, through organ culture and plant grafting we figured that gossypol was mainly synthesized in the root system. Both glanded and glandless root system had strong ability to synthesize gossypol. Our results further proved that other plant organs could synthesize the gossypol also, but their ability was weak as compared with the root systems. Previous study has shown that after biosynthesis, the gossypol was transported and stored in the pigment glands (Smith, 1962). That’s why in our study, the levels of gossypol content in glanded plants was much higher than glandless plants, as pigment glands in the former served as gossypol sink. Therefore, the absence of sink may reduce the movement of gossypol from root system to other organs or may cause the decomposition of gossypol after translocation. This speculation result may be helpful to clarify the complicated mechanism of gossypol metabolism. The technique of root and rootless plant culture may offer a new and viable way for the gene transformation. According to the previous reports on the stated compounds synthesis (Wang *et al.*, 2003; Benedict *et al.*, 2006), technique of isotope labeling and tracking could provide a more accurate way to figure out the metabolism of gossypol.

The grafting experiment in present work indicated that the root system had a notable impact on the gossypol content in the cottonseed, suggesting that the root system was the main place for gossypol biosynthesis. Besides, it was also indicated that the glanded root system might have stronger ability to synthesize gossypol than glandless ones. Furthermore, when dominant or recessive glandless plants were used as scions or rootstocks, results were almost the same. Therefore, it is suggested that the biosynthesis and transportation of gossypol in dominant and recessive plants are similar. Moreover, the effect of sunflower root system on the gossypol content of cotton scions further illustrated that the gossypol was mainly synthesized in the cotton root system. However, cotton scions were died only in a few weeks, which might have resulted from graft incompatibility. Specifically, it might have caused by the lack of vascular bundles connecting the rootstock and scion (Tiedemann, 1989). Nevertheless, the increase of gossypol content in the scions with regenerated roots, which may be induced by hormone, water and nutrient of the scions and rootstocks, helped to prove that the root system was the main organ to synthesize the gossypol.

Through gene silencing evidence, it has been proposed that gossypol biosynthesis and pigment glands formation are uncoupled. Rather they are controlled by different molecular mechanism, and restraining pigment glands formation had a feedback effect on the gossypol content (Ma *et al.*, 2016). Besides, there is a delayed gland morphogenesis trait in Australian *Gossypium* species where pigment gland formed after seed germination, and then the gossypol appears (Brubaker, 1996; Zhu and Ji, 2001). In this research, grafting changed the gossypol content, but the expression of pigment glands was unchanged, indicating that the pathways of gossypol biosynthesis and pigment glands formation were independent. The glandless root system demonstrated strong ability to synthesize gossypol; however, without pigment glands, the gossypol could not be stored. Therefore, a proper explanation could be that the gossypol was the stored material and the pigment glands were the storage tissues in cotton plants. The relationship between gossypol and pigment glands was complicated and closely linked, which need further investigation.

Earlier studied have shown that (-)-gossypol had stronger biological activity than (+)-gossypol (Puckhaber *et al.*, 2002; Wolter *et al.*, 2006; Kline *et al.*, 2008; Mellon *et al.*, 2011), and only (-)-gossypol is toxic to animals (Stipanovic *et al.*, 2006). Thus, by reducing this ratio, use of cottonseed as the feed for ruminant or non-ruminants can be increased. According to our results of organ culture, combined with the specific (+)-gossypol and (-)-gossypol contents, we speculated that the root system produced the racemic gossypol and the plant produced more (+)-gossypol which might be necessary for plant growth temporarily. The grafting between glanded cotton and sunflower further supported this phenomenon. Although these findings may help to clarify biosynthesis of the optically active gossypol, the specific roles of (+)-gossypol and (-)-gossypol in the cotton plant still need further research.

Expression profiling of the key genes participating in the gossypol biosynthesis clearly suggested that gossypol was mainly produced in the root system of glanded and glandless cotton plants, which was in accordance with the previous report (Smith, 1961). Results showed that the expression levels of key genes in gossypol biosynthesis pathway were very low in leaves, which indicated that the leaf might not have the ability to synthesize gossypol. Besides, high expression levels in the root systems of the same gene in the NILs indicated that both of the glanded and glandless cotton plants had the ability to synthesize gossypol. For the genes at the downstream of gossypol biosynthesis pathway, *GhCAD1*-*A* and *GhCYP706B1*, they all had higher expression levels in the same tissue of glanded plants than glandless plants, which also implied that the existence of pigment gland had an impact on the gossypol biosynthesis.

In this study, several new techniques which we first used to study the metabolism of gossypol can be applied to other fields. As gossypol plays a significant role in medical field as a medicine (Wang *et al.*, 1987; Coutinho, 2002; Lopez *et al.*, 2005), organ culture in vitro may offer a novel method of the gossypol production in pharmaceutical industry with a certain improvements. For the resistance research on the molecular mechanism of gossypol, when some treatments have been done to root system in vitro, the induction of responses may open the door for differential screening to discover key genes participating in gossypol pathway. Moreover, grafting technique can be used to investigate on the metabolism research.

In conclusion, our study proved that gossypol was mainly synthesized in the root systems of glanded and glandless plants. Other organs also had the ability of gossypol biosynthesis, but their ability was meager as compared to root systems. Besides, gossypol biosynthesis was not related to the expression of pigment glands, however, the presence of pigment glands did affect the gossypol content of the cotton plants.

Our findings may help to elucidate the complex network of gossypol metabolism and accelerate the process of excellent cotton breeding.

## Supplementary data

Fig. S1: The grafting combination of glanded and glandless cotton, sunflower and glanded cotton. (A) (B) The combination of glanded scion and glandless rootstock; (C) (D) The combination of glanded scion and sunflower rootstock.

Fig. S2: The processes of root tip culture and rootless seedling culture in vitro. (A) The germinated cottonseed was cut to a root tip and a rootless seedling; (B) The incubated rootless seedlings in the media; (C) The survived rootless seedlings; (D) The incubated root tips in the media; (E) The survived root systems.

Table S1: All the primers used for quantitative RT-PCR.

Table S2: The diameter and density of pigment glands in the leaves of the scions (CRI49) after grafting on different rootstocks.

Table S3: The content (mg/g) of (+), (-) and (±)-gossypol in the leaves of the scions (CRI49) after grafting on the sunflower rootstocks at different times.

Table S4: The content (mg/g) of (+), (-) and (±)-gossypol in the root systems at different times during the root tip culture in vitro.

Table S5: The content (mg/g) of (+), (-) and (±)-gossypol in the plants at different times during the rootless seedling culture in vitro

## Acknowledgements

We are grateful for the technical guidance of X. Zhang in the process of experiments. This work was funded by The National Natural Science Fund (31501342), National High Technology Research and Development Program of China (2013AA102601), The National Key Technology R&D program of China (2016YFD0101404), China Agriculture Research System (CARS-18-25), and Jiangsu Collaborative Innovation Center for Modern Crop Production.

## Reference

Adams R, Geissman TA, Edwards JD. 1960. Gossypol, a pigment of cottonseed. Chemical reviews 60, 555-574.

Bailey CA, Stipanovic RD, Ziehr MS, Huq AU, Sattar M, Kubena LF, Kim HL, Vieira RDM. 2000. Cottonseed with a high (+)- to (-)-gossypol enantiomer ratio favorable to broiler production. Journal of Agricultural and Food Chemistry 48, 5692-5695.

Benedict CR, Liu J, Stipanovic RD. 2006. The peroxidative coupling of hemigossypol to (+)- and (-)-gossypol in cottonseed extracts. Phytochemistry 67, 356-361.

Brubaker CL. 1996. Occurrence of terpenoid aldehydes and lysigenous cavities in the glandless seeds of Australian *Gossypium* species. Australian Journal of Botany 44, 601-612.

Cai Y, Xie Y, Liu J. 2010. Glandless seed and glanded plant research in cotton. Agronomy for Sustainable Development 30, 181-190.

Carrière Y, Ellers-Kirk C, Biggs R, Higginson DM, Dennehy TJ, Tabashnik BE. 2004. Effects of gossypol on fitness costs associated with resistance to Bt cotton in pink bollworm. Journal of Economic Entomology 97, 1710-1718.

Coutinho EM. 2002. Gossypol: a contraceptive for men. Contraception 65, 259-263.

Coyle T, Levante S, Shetler M, Winfield J. 1994. In vitro and in vivo cytotoxicity of gossypol against central nervous system tumor cell lines. Journal of Neuro-Oncology 19, 25-35.

Heinstein PF, Smith PF, Tove SB. 1962. Biosynthesis of C^14^-labeled gossypol. Journal of Agricultural Food Chemistry 237, 2643-2646.

Huang X, Xiao Y, Kollner TG, Zhang W, Wu J, Wu J, Guo Y, Zhang Y. 2013. Identification and characterization of (E)-beta-caryophyllene synthase and alpha/beta-pinene synthase potentially involved in constitutive and herbivore-induced terpene formation in cotton. Plant physiology and Biochemistry 73, 302-308.

Kline MP, Rajkumar SV, Timm MM, Kimlinger TK, Haug JL, Lust JA, Greipp PR, Kumar S. 2008. R-(-)-gossypol (AT-101) activates programmed cell death in multiple myeloma cells. Experimental Hematology 36, 568-76.

Kong G, Daud MK, Zhu S. 2010. Effects of pigment glands and gossypol on growth, development and insecticide-resistance of cotton bollworm (Heliothis armigera (Hübner)). Crop Protection 29, 813-819.

Leandro LL, Harlan CS, Norma LT, Thea AW. 1999. Hmg-coA reductase gene family in cotton (Gossypium hirsutum L.): Unique structural features and differential expression of hmg2 potentially associated with synthesis of specific isoprenoids in developing embryos. Plant and cell physiology 40, 750-761.

Li C, Unver T, Zhang B. 2017. A high-efficiency CRISPR/Cas9 system for targeted mutagenesis in Cotton (Gossypium hirsutum L.). Scientific Reports 7, 43902.

Liu S, Kulp SK, Sugimoto Y, Jiang J, Chang HL, Dowd MK, Wan P, Lin YC. 2002. The (-)-enantiomer of gossypol possesses higher anticancer potency than racemic gossypol in human breast cancer. Anticancer Research 22, 33–38.

Lopez LM, Grimes DA, Schulz KF. 2005. Nonhormonal drugs for contraception in men: a systematic review, Obstetrical and Gynecological Survey 60, 746–752.

Luo P, Wang Y, Wang G, Essenberg M, Chen X. 2001. Molecular cloning and functional identification of (+)-δ-cadinene-8-hydroxylase, a cytochrome P450 mono-oxygenase (CYP706B1) of cotton sesquiterpene biosynthesis. The Plant Journal 28, 95-104.

Ma D, Hu Y, Yang C, Liu B, Fang L, Wan Q, Liang W, Mei G, Wang L, Wang H, Ding L, Dong C, Pan M, Chen J, Wang S, Chen S, Cai C, Zhu X, Guan X, Zhou B, Zhu S, Wang J, Guo W, Chen X, Zhang T. 2016. Genetic basis for glandular trichome formation in cotton. Nature Communication 7, 10456.

Mao YB, Cai WJ, Wang JW, Hong GJ, Tao XY, Wang LJ, Huang YP, Chen XY. 2007. Silencing a cotton bollworm P450 monooxygenase gene by plant-mediated RNAi impairs larval tolerance of gossypol. Nature Biotechnology 25, 1307-1313.

Mellon JE, Zelaya CA, Dowd MK. 2011. Inhibitory effects of gossypol-related compounds on growth of Aspergillus flavus. Letters in Applied Microbiology 52, 406-412.

Mellon JE, Zelaya CA, Dowd MK, Beltz SB, Klich MA. 2012. Inhibitory effects of gossypol, gossypolone, and apogossypolone on a collection of economically important filamentous fungi. Journal of Agricultural Food Chemistry 60, 2740-2745.

Meng Y, Jia J, Liu C, Liang W, Heinstein P, Chen X. 1999. Coordinated accumulation of (+)-δ-cadinene synthase mRNAs and gossypol in developing seeds of *Gossypium hirsutum* and a new member of the *cad1* family from *G. arboreum*. Journal of Natural Products 62, 248-252.

Puckhaber LS, Dowd MK, Stipanovic RD, Howell CR. 2002. Toxicity of (+)- and (-)-Gossypol to the Plant Pathogen,Rhizoctonia solani. Journal of Agricultural Food Chemistry 50, 7017-7021.

Punit M, Singh ID. 1972. Growth and infestation of boll weevils on normal-glanded, glandless, and high-gossypol strains of cotton. Journal of Economic Entomology 65, 821-824.

Punit M, Singh P, Dongre AB, Narayanan SS. 1991. Variability for gossypol glands in upland cotton (Gossypium hirstum). Advance in Plant Sciences 4, 165-170.

Radloff RJ, Deck LM, Royer RE, Vander Jagt DL. 1986. Antiviral activities of gossypol and its derivatives against herpes simplex virus type II. Pharmacological Research Communications 18, 1063-1073.

Smith FH. 1961. Biosynthesis of Gossypol by Excised Cotton Roots. Nature 192, 888-889.

Smith FH. 1962. Synthesis and translocation of gossypol by cotton plant. Proceedings of the Beltwide Cotton Production Research Conference pp. 7–12.

Stipanovic RD, Bell AA, Mace ME, Howell CR. 1975. Antimicrobial terpenoids of Gossypium: 6-methoxygossypol and 6,6′-dimethoxygossypol. Phytochemistry 14, 1077–1081.

Stipanovic RD, Lopezjr, JD., Dowd MK, Puckhaber LS, Duke SE. 2006. Effect of racemic and (+)- and (–)-gossypol on the survival and development of *Helicoverpa zea* larvae. Journal of Chemistry Ecology 32, 959-968.

Stipanovic RD, Lopezjr JD, Dowd MK, Puckhaber LS. Duke SE. 2008. Effect of racemic, (+)- and (-)-gossypol on survival and development of *Heliothis virescens* larvae. Environmental Entomology 37, 1081–1085.

Sunilkumar G, Campbell LM, Puckhaber L, Stipanovic RD, Rathore KS. 2006. Engineering cottonseed for use in human nutrition by tissue-specific reduction of toxic gossypol. Proceeding of the National Academy of Sciences of the United States of America 103, 18054-18059.

Tegos G, Stermitz FR, Lomovskaya O, Lewis K. 2002. Multidrug pump inhibitors uncover remarkable activity of plant antimicrobials. Antimicrobial Agents and Chemotherapy 46, 3133-3141.

Tiedemann R. 1989. Graft union development and symplastic phloem contact in the heterograft Cucumis sativus on Cucurbita ficifolia. Journal of Plant Physiology 134, 427-440.

Townsend BJ, Poole A, Blake CJ, Llewellyn DJ. 2005. Antisense suppression of a (+)-delta-cadinene synthase gene in cotton prevents the induction of this defense response gene during bacterial blight infection but not its constitutive expression. Plant Physiology. 138, 516-528.

Wang JY, Cai Y, Gou JY, Mao YB, Xu YH, Jiang WH, Chen XY. 2004. VdNEP, an elicitor from *Verticillium dahliae*, induces cotton plant wilting. Applied and Environmental Microbiology 70, 4989-4995.

Wang N, Zhou L, Guan M, Lei H. 1987. Effect of (-) and (+) gossypol on fertility in male rats. Journal of Ethnopharmacology 20, 21-24.

Wang Y, Davila-Huerta G, Essenberg M. 2003. 8-Hydroxy-(+)-δ-cadinene is a precursor to hemigossypol in *Gossypium hirsutum*. Phytochemistry 63, 219-225.

William JL, Ellers-Kirk C, Orth RG, Gassmann AJ, Head G, Tabashnik BE, Carrière Y. 2011. Fitness cost of resistance to Bt cotton linked with increased gossypol content in pink bollworm larvae. PLoS One 6, e21863.

Wolter KG, Wang SJ, Henson BS, Wang S, Griffith KA, Kumar B, Chen J, Carey TE, Bradford CR, D’Silva NJ. 2006. (-)-Gossypol inhibits growth and promotes apoptosis of human head and neck squamous cell carcinoma in vivo. Neoplasia 8, 163-172.

Xiang S, Yang W. 1993. Studies on gossypol and its enantiomers in the seeds of cotton Gossypium. Scientia Agricultural Sinica 26, 31-35.

Xu YH, Wang JW, Wang S, Wang JY, Chen XY. 2004. Characterization of GaWRKY1, a cotton transcription factor that regulates the sesquiterpene synthase gene (+)-delta-cadinene synthase-A. Plant physiology 135, 507-515.

Ye W, Chang H, Wang L, Huang Y, Shu S, Dowd MK, Wan PJ, Sugimoto Y, Lin Y. 2007. Modulation of multidrug resistance gene expression in human breast cancer cells by (-)-gossypol-enriched cotton oil. Anticancer Research 27, 107-116

Zhou M, Zhang C, Wu Y, Tang Y. 2013. Metabolic engineering of gossypol in cotton. Applied microbiology and biotechnology 97, 6159-6165.

Zhu S, Ji D. 2001. Inheritance of the delayed gland morphogenesis trait in Australian wild species of *Gossypium*. Chinese Science Bulletin 46, 1168-1174.

